# Modelling prodromal Parkinson’s disease: gastrointestinal pathology and early motor deficits in mice after transient rotenone exposure

**DOI:** 10.64898/2026.10.23.761386

**Authors:** Sharmelee Selvaraji, Jasmine Mosberger, Karen K.L. Pang, Gari L. Eberly, Ye Ji Kim, Rajib Mondal, Rebecca Leomi, Polina Anikeeva

**Affiliations:** McGovern Institute for Brain Research, Massachusetts Institute of Technology, Cambridge, MA, USA; K. Lisa Yang Brain-Body Center, Massachusetts Institute of Technology, Cambridge, MA, USA; Research Laboratory of Electronics, Massachusetts Institute of Technology, Cambridge, MA, USA; Neuroscience Program, Wellesley College, Wellesley, MA, USA; Department of Brain and Cognitive Sciences, Massachusetts Institute of Technology, Cambridge, MA, USA; Harvard-MIT Division of Health Sciences and Technology, Massachusetts Institute of Technology, Cambridge, MA, USA; Department of Materials Science and Engineering, Massachusetts Institute of Technology, Cambridge, MA, USA

## Abstract

Early gastrointestinal (GI) dysfunction is a hallmark feature of Parkinson’s disease (PD), yet models capturing prodromal gut and brain pathology in PD remain limited. Motivated by the increased PD risk associated with pesticide exposure, we monitored motor behavior, brain pathology, and GI structure and function over 9 months following 5 daily doses of rotenone in mice. Transient rotenone exposure induced significant nigrostriatal pathology, ∼48% loss of dopaminergic neurons and increased α-synuclein aggregation in the substantia nigra, without significant changes in gross motor function. Notably, we identified pronounced GI dysfunction including reduced colonic transit velocity, altered motility dynamics, and delayed whole-gut transit. These functional changes were accompanied by enteric dopaminergic neurodegeneration, increased colonic α-synuclein accumulation, increased colonic crypt depth, and goblet cell hyperplasia with elevated mucin content per goblet cell. The findings are consistent with prodromal PD, suggesting the utility of this model for investigating early disease mechanisms and intervention strategies.

## Introduction

Parkinson’s disease (PD) is the fastest growing neurodegenerative motor disorder, affecting over 10 million people worldwide (https://www.parkinson.org/understanding-parkinsons/statistics), with 90% of PD cases being sporadic in nature^1^. PD is characterized by the loss of dopaminergic (DA) neurons and pathological aggregation of α-synuclein protein in the substantia nigra pars compacta (SNpc), manifesting in impaired voluntary motor function^2^. However, PD pathology is not confined to the brain, and over 80% of patients experience gastrointestinal (GI) symptoms such as constipation years before motor impairments develop^3,4^. These early non-motor features motivate the investigation of the bidirectional gut-brain communication in PD pathogenesis^5^. Accordingly, competing brain-first and gut-first models have emerged in the past two decades. The brain-first model suggests that PD pathology originates in the central nervous system (CNS) and the neurodegeneration secondarily impair enteric (gut) function^6^. The gut-first model proposes that some PD pathologies may originate in the enteric nervous system (ENS) following environmental insults to the GI tract before spreading to the CNS via vagal or dorsal root pathways (Braak’s hypothesis)^7–11^. Notably, α-synuclein inclusions have also been identified in enteric neurons of post-mortem PD patients^12^. Despite the controversies surrounding the gut-first hypothesis, over the past two decades, a growing body of clinical, epidemiological and experimental evidence has strengthened support for Braak’s hypothesis, including the delineation of staging patterns for α-synuclein pathology, associations between vagotomies and reduced PD risk, and preclinical studies demonstrating gut-to-brain transmission of α-synuclein^7,13–15^. These studies underscore the need for mechanistic studies of GI pathology in PD as they may accelerate the development of early diagnostics and lead to improved quality of life for PD patients.

Despite the growing recognition of the GI pathology in PD, the most reliable experimental PD models remain restricted to the brain, relying on acute neurotoxin exposure or direct central α-synuclein overexpression in the SNpc^16^. These models do not reproduce early GI dysfunction or α-synuclein accumulation in the ENS observed in patients. To recapitulate ENS pathology, pre-formed α-synuclein fibrils have been directly injected into the GI tract in rodent models^15,17^. These studies observed propagation of α-synuclein pathology from the gut to the brain, demonstrating DA degeneration in the SNpc and motor dysfunction, however they did not directly model the mechanisms underlying the pathological α-synuclein fibril formation.

Clinical data linking exposure to herbicides and insecticides to the increased risk of PD in agricultural communities^18–21^ motivated the development of environmental animal models of PD with the goal of recapitulating the progressive nature of the disease^22–24^. For instance, exposure to rotenone, a naturally derived insecticide and piscicide which inhibits mitochondrial complex I, has been shown to cause neuropathological deficits that closely resemble PD in humans and mice^22,25,26^. However, the effects of pesticide exposure on GI function as well as the staging of the gut and brain pathology in these environmental models of PD are yet to be fully elucidated.

Here, we used transient rotenone exposure as a model to investigate early gut pathology and the progression of central neurodegeneration. In this environmental model of PD, 7-month-old mice were intraperitoneally injected with rotenone for 5 consecutive days and then monitored for 9 months to capture progressive aspects of pathology. The effects of rotenone exposure were evaluated via functional gut assays, immunohistochemical analyses, and motor behavior assessments. Consistent with prior observations, we found that environmental exposure to rotenone resulted in α-synuclein aggregation and loss of DA neurons in SNpc. Notably, our data revealed enteric α-synuclein aggregation and significant pathological changes in gut morphology and function, while motor alterations were subtle and consistent with the prodromal phase of PD. Together, these findings suggest that transient rotenone exposure offers a physiologically grounded platform for studying early gut and brain pathology in PD.

## Results

### Effects of rotenone exposure on DA neuron viability and α-synuclein accumulation in the SNpc

Adult male mice (7 months old) were dosed with rotenone (2.5mg/kg) or vehicle (2% DMSO; 98% Miglyol 812) via intraperitoneal injection for 5 consecutive days and were kept on observation for 9 months^26^ (**Fig. 1a**). Male mice were used in this study since sporadic PD is more prevalent in males^27^. The body weights of these mice were monitored weekly, and the increase in body weight was not significantly different between the rotenone (42.25 ± 4.80 g) and vehicle (42.56 ±5.16 g) groups at the end of the study (**Fig. 1b, c**).

**Figure 1.**
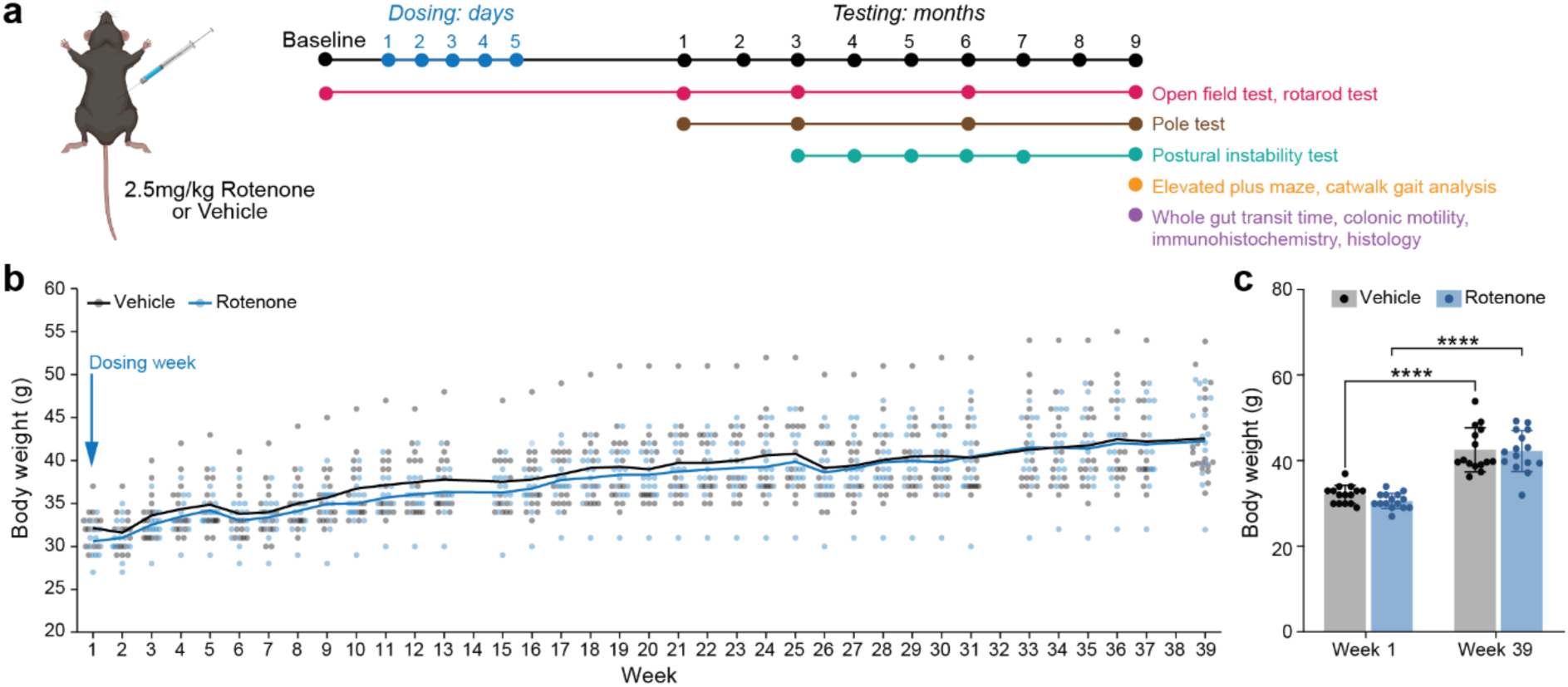
Transient rotenone exposure does not alter body weight over a 9-month period. **(a)** Experimental timeline. 7-month-old adult male mice received intraperitoneal injections of rotenone or vehicle for 5 consecutive days and were monitored for 9 months. A battery of behavioral assays was conducted during the 9 months followed by functional assays of gut transit and histology. **(b)** Longitudinal body weight measurements recorded weekly over the 9-month observation period. **(c)** Body weights at the start (Week 1) and end (Week 39) of the 9-month study period. Significant increases in body weight were observed in both vehicle (****p<0.0001) and rotenone (****p<0.001) mice between the start and end of the study period respectively. No statistical significance was observed between rotenone- and vehicle-injected mice. Statistical analysis was performed using unpaired t-test with Welch’s correction. Data are presented as mean ± standard deviation (SD); n =15 mice per group. Image of mouse was created with BioRender.

To assess the impact of rotenone exposure on the nigrostriatal DA system, DA neuronal integrity was examined in the SNpc at the conclusion of the 9 months period following the 5-day dosing protocol. Immunofluorescence analysis revealed a 47.7% (95% CI: –262.7 to – 83.78) reduction in tyrosine-hydroxylase expressing (TH^+^) DA neurons in the SNpc of rotenone-injected mice as compared to vehicle-injected mice (**Fig. 2a, b**). Aggregation of α-synuclein was observed in rotenone-injected mice through Avidin-Biotin Complex (ABC) staining for total α-synuclein, although the increase was not statistically significant (**Fig. 2c, d**).

**Figure 2.**
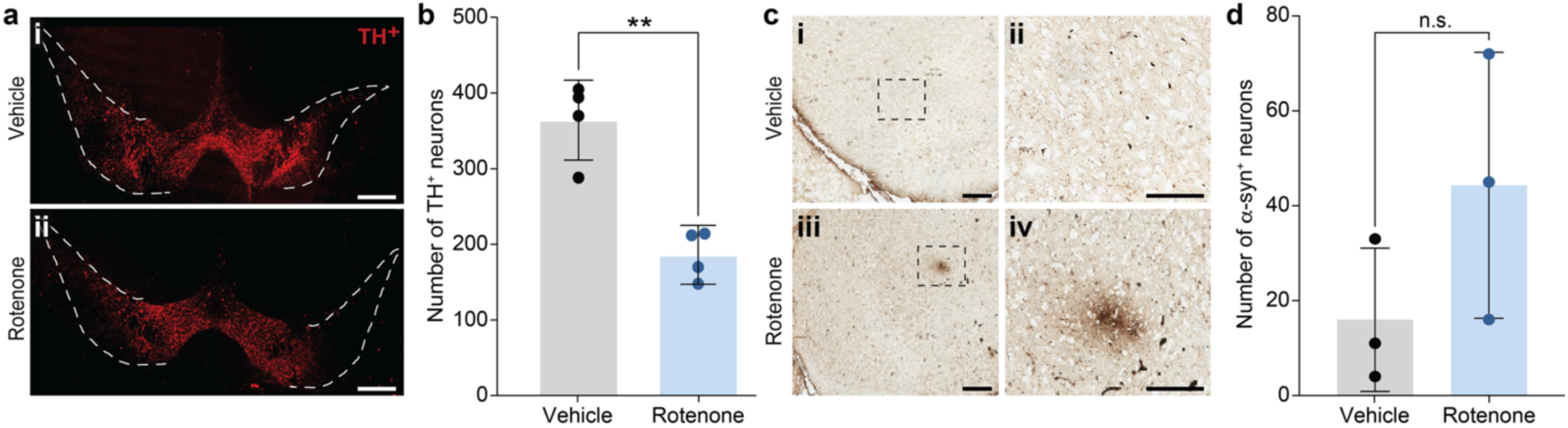
Transient systemic rotenone exposure induces dopaminergic neuronal loss and α-synuclein pathology in the mouse brain. **(a)** Representative immunofluorescence images of tyrosine hydroxylase (TH)-positive neurons in the SNpc from **(i)** vehicle- and **(ii)** rotenone-treated mice 9 months after the 5-day exposure. Scale bars, 200 µm. **(b)** Number of TH^+^ neurons in the SNpc in rotenone-treated mice and vehicle controls (Mann-Whitney U test, **p*=*0.0148). **(c)** Representative images of total α-synuclein immunostaining in the SNpc of (i, ii) vehicle control and (iii, iv) rotenone-treated mice. Rotenone-injected mice exhibited punctate and clustered α-synuclein inclusions, whereas vehicle-injected mice showed diffuse α-synuclein staining without aggregates. (i, iii) Scale bars, 200 µm. (ii, iv) Scale bars, 100 µm. **(d)** α-synuclein immunoreactivity, showing increased accumulation in rotenone-injected mice compared to controls, which did not reach statistical significance (Mann-Whitney U test, p=0.200; ns, not significant). Data are presented as mean ± SD; n=3-5 mice per group.

### Longitudinal effects of transient rotenone exposure on motor function

Motor symptoms are the key diagnostic features of PD^28,29^. Consequently, we performed motor assessments in mice exposed to rotenone or vehicle for 5 days at multiple time points over the 9-month monitoring period. To assess bradykinesia and overall locomotor function, a pole test was performed at 1, 3, 6, and 9 months, but no significant difference was observed between groups or timepoints for the time taken to turn 180° (**Fig. 3a**) or the latency to climb down the pole (**Fig. 3b**). To assess gross locomotor activity and potential anxiety-like behaviors, the open field test (OFT) was performed at baseline, prior to rotenone (or vehicle) exposure, and then repeated 1, 3, 6 and 9 months following the 5-day injection protocol. The time spent by the mice in the periphery was not significantly different between the rotenone and vehicle groups at baseline, 1, 3 and 6 months post-treatment (**Fig. 3c**). However, at 9 months post-treatment, rotenone-injected mice spent a significantly shorter time in the periphery compared to vehicle-injected mice, with the total distance travelled by the mice remaining comparable between groups and across timepoints (**Fig. 3d)**. The time immobile in the OFT setup was significantly lower in the rotenone-injected mice as well **(Fig. 3e)**. Contrary to expectation, these indicate altered exploratory behaviors, consistent with reduced anxiety-like behavior, without a corresponding increase in overall locomotor activity. Motor coordination, endurance and balance were further evaluated via a rotarod assay at 1, 3, 6 and 9 months post-exposure. Surprisingly, the latency to fall was longer in rotenone-injected mice compared to vehicle controls at 6 months post-exposure, and a similar but not statistically significant trend was observed at 9 months (**Fig. 3f**). To probe whether the shorter time spent in the periphery of the open field and the increased latency of first fall from the rotarod of rotenone-injected mice were due to anxiety-influenced locomotion, the elevated plus maze assay was performed. Contrarily, rotenone-injected mice spent significantly lesser time in the closed arms than vehicle-injected mice implying reduced anxiety-like behavior rather than increased anxiety (**Fig. 3g**). These results reveal that, unlike models employing chemical depletion of DA neurons in the brain^30^, transient rotenone exposure does not cause overt PD-like motor deficits or impaired coordination over the 9-month period. The lack of significant gross motor impairment is consistent with the <50% SNpc DA neuron depletion, as prior studies have shown that observable gross motor impairments emerge only after more than 70% DA depletion^31^.

**Figure 3.**
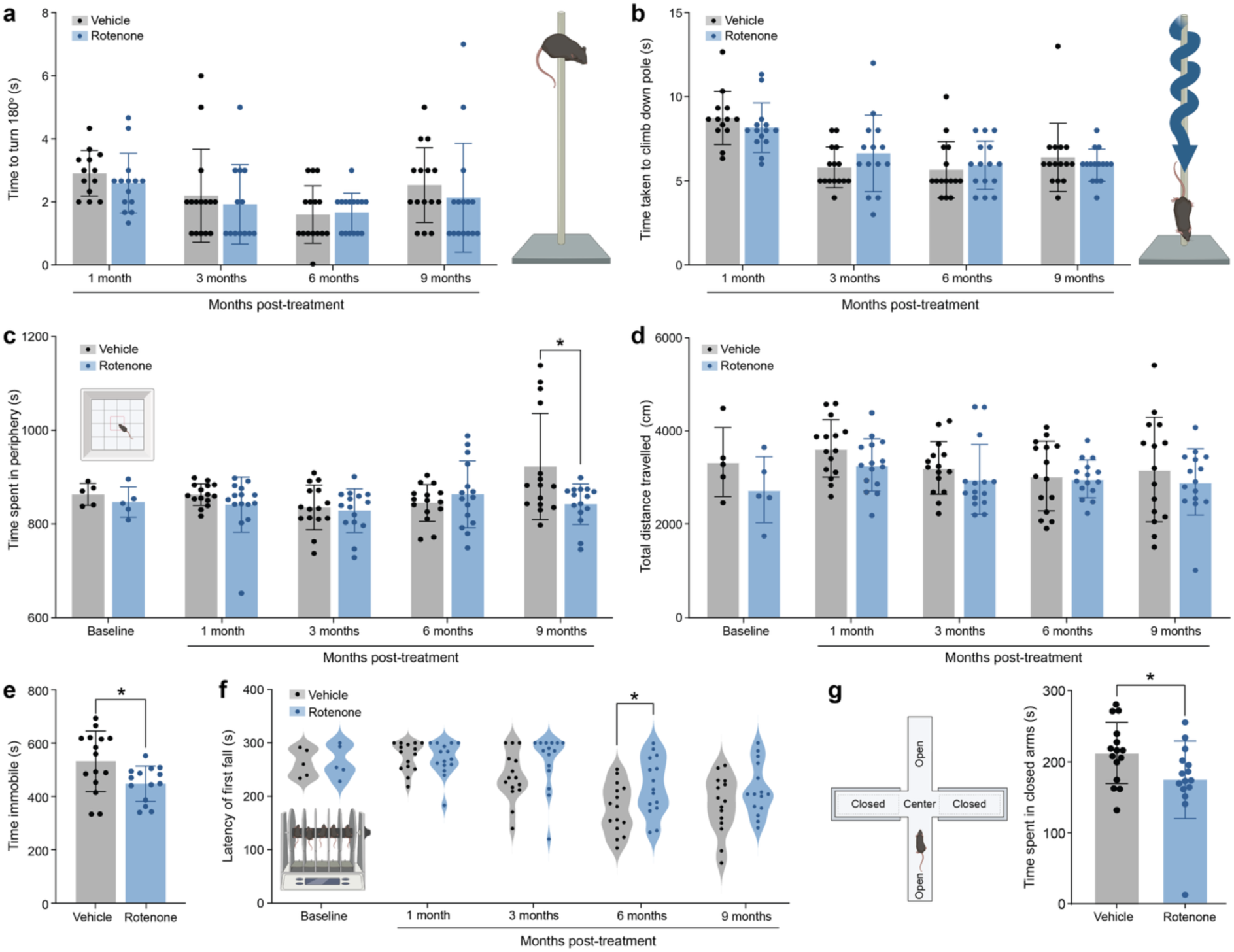
Transient rotenone exposure does not induce major motor deficits in mice over a 9-month period. **(a–b)** Pole test assessing bradykinesia and motor coordination. (**a**) No significant differences were observed between rotenone- and vehicle-injected mice in the time to turn 180° or (**b**) the latency to descend the pole at 1, 3, 6, and 9 months post-treatment. **(c–e)** Open field test (OFT) evaluating locomotor activity and anxiety-like behavior. (**c**) Time spent in the periphery and (**d**) total distance travelled did not differ between groups at most time points except at 9 months for time spent in periphery (Tukey’s multiple comparisons test, *p=0.0195). **(e)** At 9 months post-treatment, the duration of immobile periods was compared, with a shorter immobile period in rotenone mice (unpaired t-test with Welch’s corrections, *p=0.0224) observed. **(f)** Rotarod assay measuring motor coordination and balance. Latency to first fall was significantly longer in rotenone-injected mice at 6 months (Tukey’s multiple comparisons test, *p=0.0310). **(g)** Elevated plus maze assessing anxiety-related locomotion. Rotenone-injected mice spent significantly less time in the closed arms than vehicle-injected mice (unpaired two-tailed test, *p=0.0453). Data are presented as mean ± SD; *n* = 15 mice per group. Images of the pole test, open field set-up, rotarod and EPM were created in BioRender.

Although no significant changes in gross motor coordination were observed, we sought to assess the effects of transient rotenone exposure on fine motor coordination. To assess postural reflex impairment and balance control, the postural instability test (PIT) was conducted. Both rotenone- and vehicle-treated groups showed a significant increase in the distance to trigger compensatory step at 9 months as compared to 3 months (**Fig. 4a**), suggesting learned balance control on the PIT with repeated exposure. Nonetheless, the gradient of increase was steeper in the vehicle-injected mice compared to the rotenone-injected mice. Notably, rotenone-injected mice displayed a shorter distance to trigger compensatory step compared to vehicle-injected mice at 4, 6 and 9-month timepoints respectively, indicating impaired postural reflex and balance. The number of steps taken for the same distance covered were significantly lower 4 months post-treatment in rotenone-injected mice compared to vehicle-injected mice, indicating an initial decline in postural stability with rotenone exposure (**Fig. 4b**).

**Figure 4.**
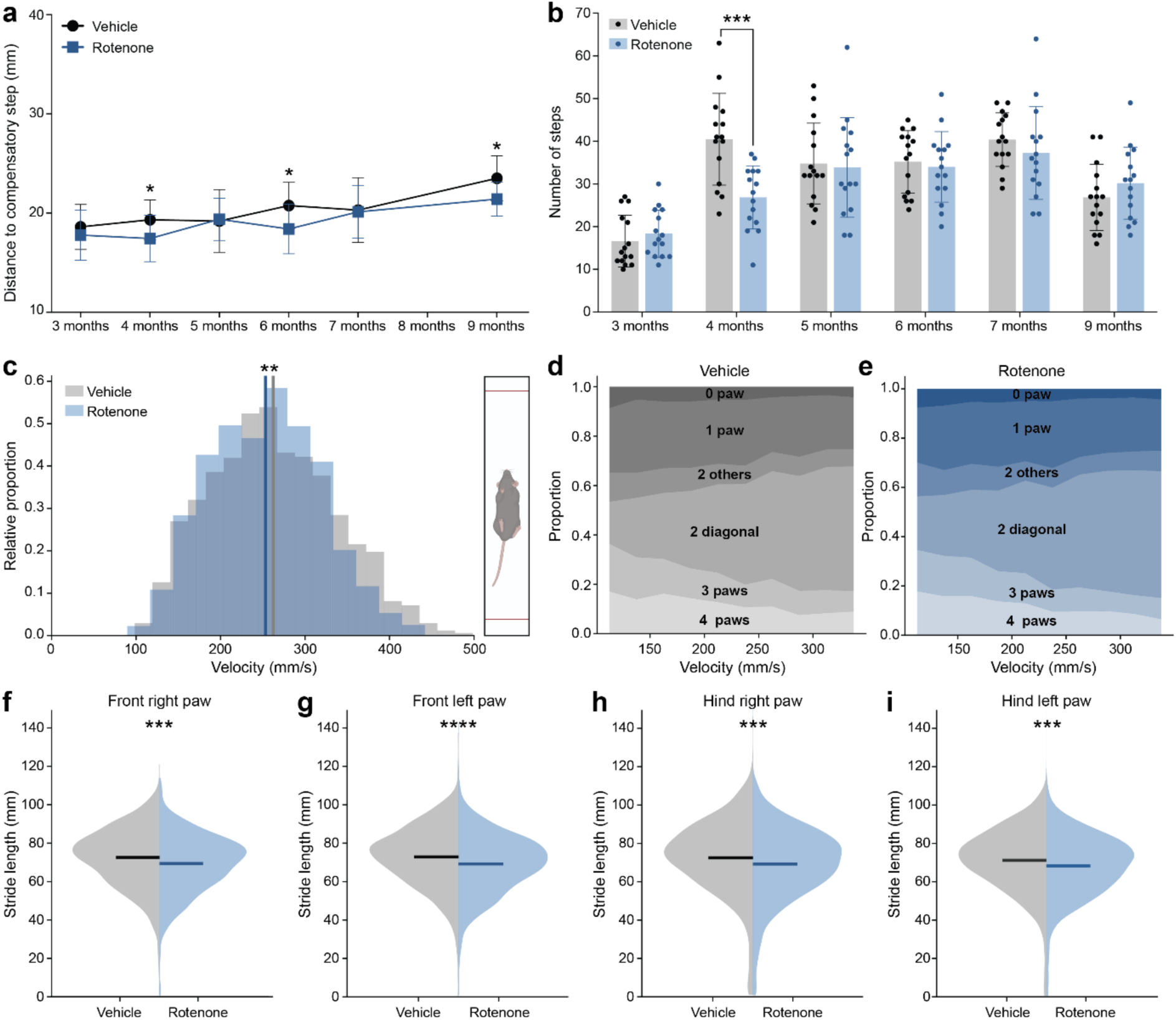
Transient rotenone exposure impairs fine motor coordination and balance in mice. **(a–b)** Postural instability test assessing balance control and postural reflex. **(a)** Both rotenone- and vehicle-injected mice showed increased distance to trigger compensatory step over 9 months, indicative of learned balance control. However, rotenone-injected mice exhibited shorter distances at 4 (Tukey’s multiple comparison test, *p=0.0261), 6 (Tukey’s multiple comparison test, *p=0.0126), and 9 (Tukey’s multiple comparison test, *p=0.0081) months compared to vehicle controls, suggesting impaired postural reflex. **(b)** The number of steps taken for the same distance was significantly lower at 4 months (Tukey’s multiple comparison test, *p=0.0005) in rotenone-injected mice. **(c–j)** CatWalk gait analysis evaluating locomotion and gait patterns. **(c)** Movement velocity was significantly reduced in rotenone-injected mice at 9 months post-treatment (Rank sum test, **p =0.00213). **(d–e)** Diagonal support measures showed no notable differences, with only subtle variations at low velocities. **(f)** Stride length analysis of front right paw (***p=6.084 x 10^-10^), front left paw (****p=4.797 x 10^-12^), hind right paw (***p=9.598 x 10^-7^) and hind left paw (***p=2.752 x 10^-7^) revealed consistently reduced stride length in all four paws. Data are presented as mean ± SD; n = 15 mice per group.

Gait disruption is a hallmark feature of PD in patients^32^, and consequently gait features were assessed at 9 months following a 5-day exposure to rotenone (or vehicle) using a catwalk assay. An automated analysis pipeline, GaitPattern^33^, was then employed to compare gait features between the groups. The movement velocity was significantly lower in the rotenone-injected mice compared to the vehicle-injected controls (**Fig. 4c**). The variations in diagonal support, which assesses the gait pattern where the bodyweight is supported by two diagonally opposite limbs during quadrupedal locomotion, showed only subtle differences in the low velocity range (**Fig. 4d,e**). However, the stride length across all four individual paws of rotenone-injected mice was significantly shorter than that of vehicle-injected mice (**Fig. 4f-i**).

### Effects of transient rotenone exposure on colonic motility

Given that constipation is a common comorbidity of PD (80% of cases) and often predates motor deficits^3,4^, we sought to investigate the impact of transient rotenone exposure on GI motility. First, to assess global changes in the GI motility, at 9 months following the 5-day exposure to rotenone (or vehicle), mice were orally gavaged with carmine dye (6% w/v in 0.5% methylcellulose solution) and the time taken to expel the first dyed fecal pellet was recorded.

Rotenone-injected mice took a longer time to pass the first dyed fecal pellet than vehicle-injected mice, indicating a trend towards a slower whole-GI motility, but this change was not statistically significant (**Fig. 5a**). Transit velocity, approximated by dividing colon length by whole gut transit time, was in fact significantly lower in the rotenone group as compared to controls (**Fig. 5b**), which is consistent with the trend for reduced colon length in these mice^34,35^ (**Fig. 5c,d**).

**Figure 5.**
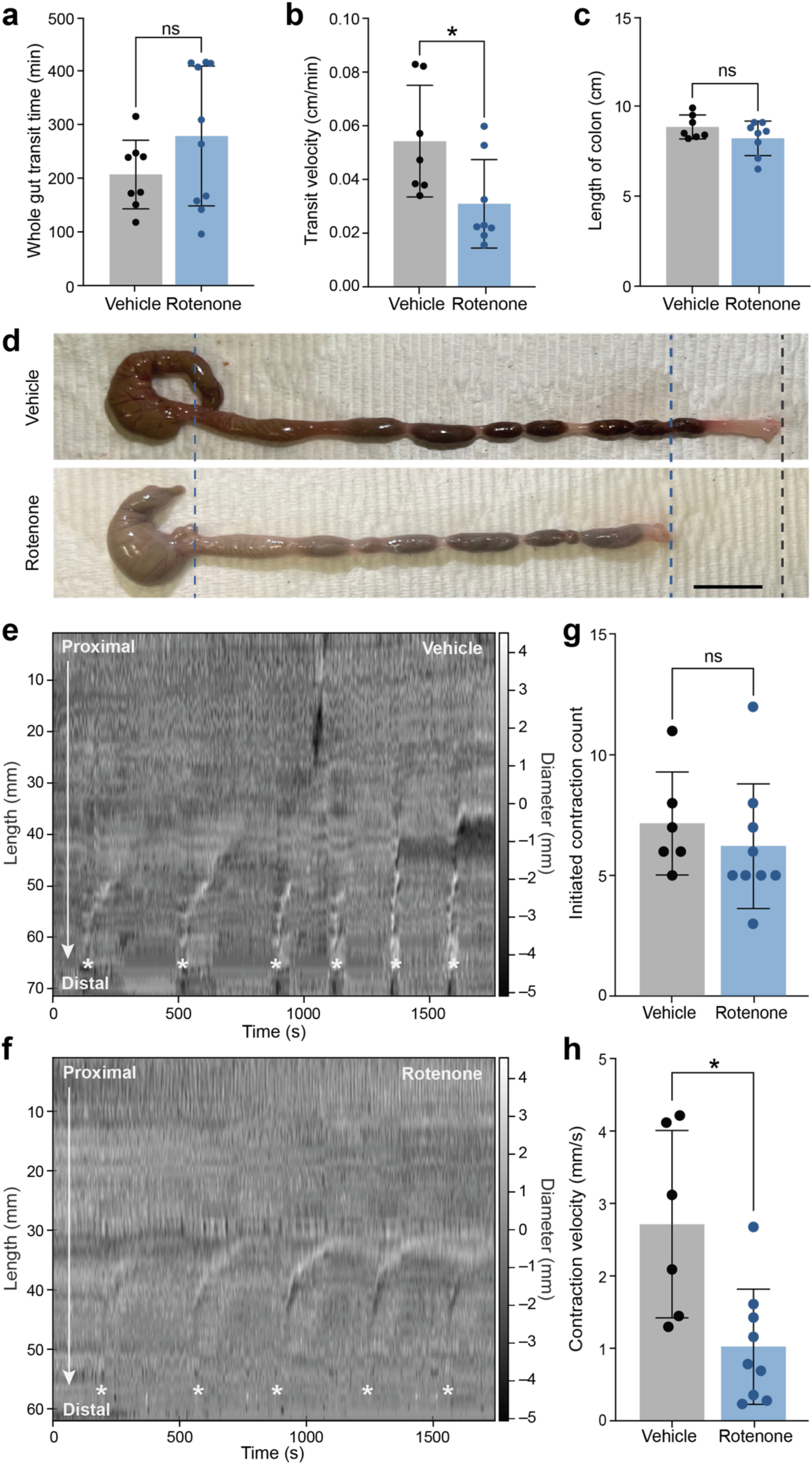
Transient rotenone exposure alters colonic transit velocity and contraction dynamics. **(a)** Whole-gut motility assessed by carmine dye gavage. Rotenone-injected mice exhibited a longer time to expel first dyed fecal pellet compared to vehicle-injected mice, though this difference was not statistically significant (Mann-Whitney U test, p=0.360). **(b)** Transit velocity, normalized to colon length, was significantly slower in rotenone-injected mice compared to vehicle-injected mice (Mann-Whitney U test, *p=0.0289), reflecting combined effects of slower whole-gut transit and shorter colon length. **(c)** Colon length was shorter in rotenone-injected mice, though not statistically significant (Mann-Whitney U test, p=0.412). **(d)** Representative images of vehicle- and rotenone-injected mouse colons after 9-month study period. **(e, f)** Representative spatiotemporal maps of *ex-vivo* colonic motility used to quantify contraction dynamics. White asterisk (*) indicates each colonic contraction. **(g, h)** Quantification of contraction features in the proximal colon: **(g)** number of contractions initiated was not significantly different between the two groups (unpaired t-test with Welch’s correction, p=0.4560) but **(h)** contraction velocity was significantly slower in rotenone-injected mice (unpaired t-test with Welch’s correction, *p=0.0224). Data are presented as mean ± SD; *n* = 6-10 mice per group; ns, not significant Scale bar, 1cm.

To further probe intrinsic colonic motility, an *ex vivo* motility assay was used to map spatiotemporal representations of colonic contractions (**Fig. 5e,f**). Although the number of initiated contractions (**Fig. 5g**) in the proximal colon were not significantly different between the rotenone and control groups, the contraction velocity was significantly slower in the rotenone group (**Fig. 5h**).

### Effects of transient rotenone exposure on structural and cellular architecture of the colon

The observed reduction in colonic transit velocity following rotenone exposure motivated further assessment of colon anatomy and cytoarchitecture. Given that the myenteric plexus that regulates GI motility harbors DA neurons^36^, we tested whether these neurons undergo degeneration upon exposure to rotenone akin to DA neurons in the brain. Rotenone-injected mice showed a statistically significant decline in TH^+^ neurons in the myenteric plexus in the colon compared to vehicle-injected mice (**Fig. 6a-c**). Furthermore, akin to the findings in the SNpc, immunostaining for α-synuclein revealed significant accumulation and prominent aggregate-like inclusions in the colon of rotenone-injected mice, while no notable aggregation was observed in vehicle-injected controls (**Fig. 6d-f**). Structural analysis with Hematoxylin and Eosin (H&E) staining revealed that rotenone-injected mice had significantly longer colonic crypt depth compared to vehicle-injected controls, suggesting an inflammatory phenotype^37,38^ (**Fig. 6g,h**). Finally, Alcian blue staining, which marks acidic mucins in goblet cells lining the colon, showed significantly higher concentration of mucin content per goblet cell and a significantly higher number of goblet cells in rotenone-injected mice compared to vehicle-injected controls, indicating compromised mucus barrier integrity in the former^39,40^ (**Fig. 6i,j**).

**Figure 6.**
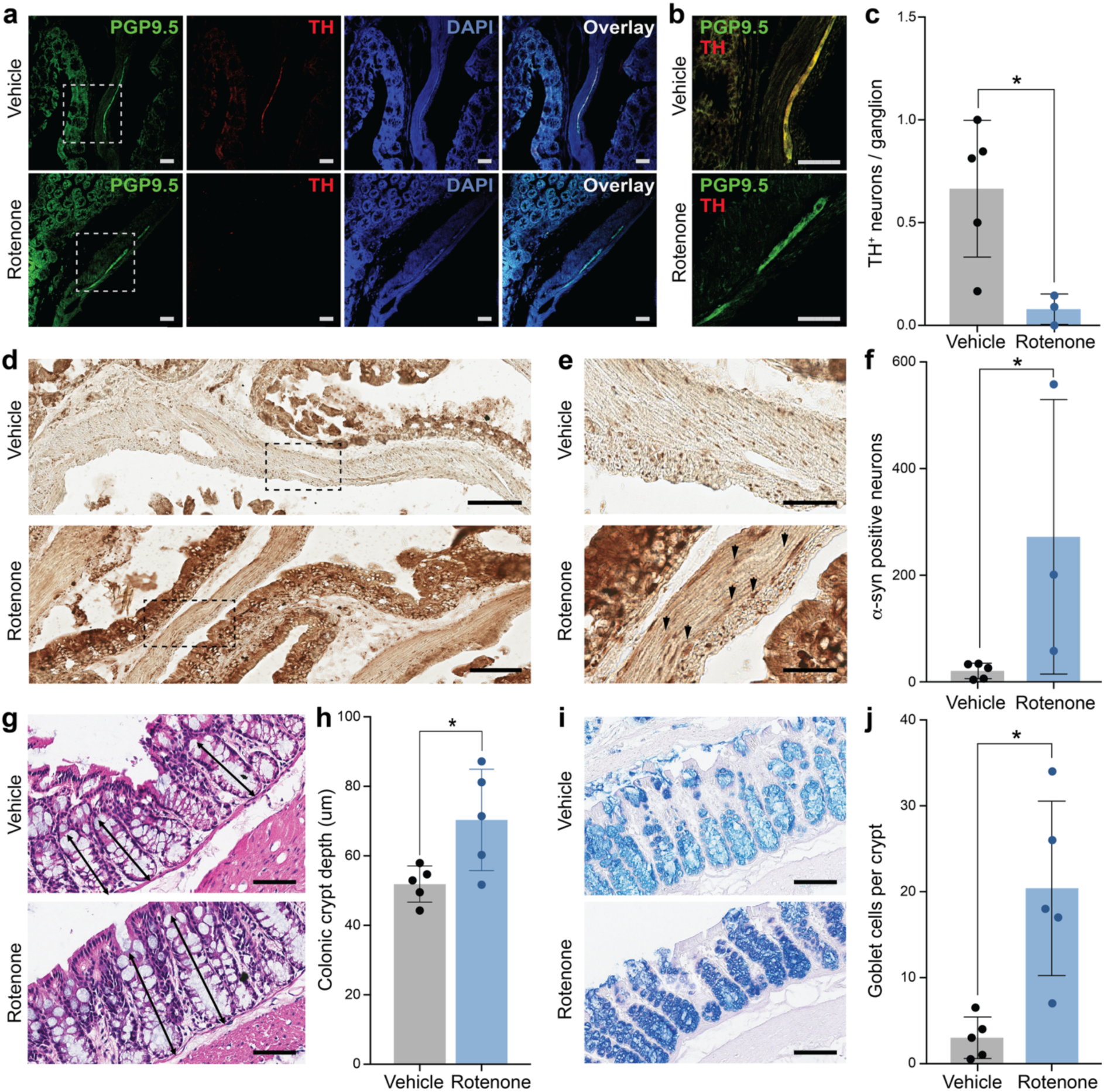
Transient rotenone exposure induces enteric dopaminergic neuronal loss, α-synuclein aggregation, altered crypt architecture, and impaired mucus barrier in the mouse colon. **(a–b)** Immunohistochemical analysis of tyrosine hydroxylase -positive (TH^+^) neurons in the myenteric plexus of the colon. Scale bars, 50 µm. **(c)** Rotenone-injected mice showed a significant reduction (Mann-Whitney U test, *p=0.0357) in TH^+^ neurons compared to vehicle-injected controls, indicating enteric dopaminergic neuronal loss. **(d-e)** Avidin-Biotin Complex (ABC) staining of α-synuclein in the colon. Scale bars, 50 µm **(d),** 100 µm **(e). (f)** Rotenone-injected mice exhibited significant increase in α-synuclein accumulation and aggregate-like inclusions relative to vehicle-injected mice, (Mann-Whitney U test, *p= 0.0357). **(g)** Hematoxylin and Eosin (H&E) staining demonstrating structural changes in the colon. **(h)** Rotenone-injected mice displayed significantly longer (unpaired t-test with Welch’s correction, *p=0.0446) colonic crypts compared to vehicle-injected mice, suggesting altered tissue architecture. **(i)** Alcian blue staining of goblet cells assessing mucus barrier function. **(j)** Rotenone-injected mice showed significantly higher mucin content per goblet cell and an increased number of goblet cells relative to vehicle-injected mice (unpaired t-test with Welch’s correction, *p=0.0168), indicative of compromised mucus barrier integrity. Scale bars, 100 µm **(g, i)**. Data are presented as mean ± SD; *n* = 3-5 mice per group.

## Discussion

Rotenone is widely used to induce PD-like phenotype in animal models^41^. While gastroparesis and constipation have been previously observed in the rotenone models^42^, capturing pre-motor enteric pathology and dysmotility in isolation has proven challenging, limiting the utility of these environmental models for studies of gut-brain interactions in PD. Van Laar and colleagues previously showed that a transient 5-day exposure to rotenone elicited PD-like motor deficits and DA neuron degeneration in the brain even after 9 months^26^. However, the effects of this treatment regime on the gut remained to be explored.

Here, we demonstrate that the 5-day exposure to rotenone is sufficient to induce long-lasting gut and brain pathology in mice. By exhibiting a modest motor phenotype, but significant (∼48%) DA neuron degeneration in the SNpc, and profound changes in the gut transit, morphology, and cytoarchitecture, this model mimics key features of the prodromal phase of PD pathology^43,44^. Restricting the length of rotenone exposure enabled observation of pathological and functional changes in the absence of overt systemic toxicity. Moreover, given the toxicity of rotenone, the changes observed in this transient model can be attributed to pathological changes to physiology rather than prolonged sickness behavior^25^.

While 5-day rotenone exposure had no significant effect on gross motor performance, rotenone-treated mice exhibited some unexpected behavior. Six months following treatment, these mice had a significantly higher latency of first fall on the rotarod compared to vehicle-injected controls. The treated mice also showed reduced anxiety-like behavior in the open field and elevated plus maze assays. These unexpected observations could, at least in part, be due to potential compensatory effects of α-synuclein aggregation on emotional regulation in PD mice^45,46^, which in turn could affect performance on the rotarod and other behavioral assays.

In contrast to gross motor assays, quantitative analysis of fine motor coordination revealed subtle but consistent impairments in rotenone-treated mice. The automated PIT analysis quantified step distance using successive forelimb placements rather than nose displacement, providing an objective and scalable measure of postural control. The observed changes in both step distance and step number in rotenone-treated mice over time suggest dynamic adaptations in motor output that may reflect early dysfunction in basal ganglia and brainstem circuits involved in balance and postural regulation. Consistent with these findings, gait analysis using a catwalk assay showed reduced movement velocity and shortened stride length across all four paws in rotenone-injected mice. This suggests intact gross coordination but impaired stride scaling and fine motor control. Such pattern closely resembles early gait abnormalities observed in PD patients like reduced stride length and shuffling gait, which often precedes more overt motor symptoms^47^. Together, these findings suggest that transient rotenone exposure leads to subclinical motor deficits that are detectable through quantitative analyses of fine motor coordination.

Unlike the subtle motor deficits, GI dysmotility was a prominent and persistent feature of the rotenone-injected mice, aligned with the clinical presentations of constipation prevalent in early-stage PD patients^48–50^. The functional changes were accompanied by distinct cellular and structural abnormalities in the colon. Rotenone-injected mice displayed significant loss of TH^+^ neurons in the myenteric plexus, mirroring the vulnerability of dopaminergic neurons in the brain. α-synuclein aggregation was also markedly increased in the colon, suggesting that pathological protein accumulation is not restricted to the central nervous system in this model. Structurally, increased crypt depth and alterations in the goblet cell number and mucin content were indicative of disrupted epithelial homeostasis and impaired mucus barrier function^51,52^. Therefore, the combination of enteric neurodegeneration, α-synuclein aggregation and epithelial dysfunction provides a link between environmental toxin exposure and GI pathology relevant to early-stage PD.

While this study provides evidence for gut and brain pathology following transient rotenone exposure, several limitations should be acknowledged. Despite robustly capturing the prodromal phase of PD, the 9-months duration of this model limits its utility in studies seeking to evaluate multiple diagnostic or therapeutic hypothesis. Recently, oral gavage of low dose rotenone for up to 10 weeks was shown to induce constipation phenotype in mice^35^, inviting further functional characterization in this accelerated model of GI pathology. Additionally, while we observed α-synuclein aggregation in the brain and colon, the temporal sequence and potential propagation pathways remain to be elucidated. Future studies incorporating longitudinal sampling and vagotomy experiments would help clarify the mechanisms linking gut pathology to central neurodegeneration.

The transient rotenone exposure model demonstrates how a brief period of environmental insult initiates a cascade of long-lasting pathological changes affecting the gut and brain physiology, manifesting in GI dysfunction preceding motor impairments. This temporal pattern closely resembles the prodromal phase of PD, during which non-motor symptoms such as constipation often appear decades before diagnosis in PD patients^48,50^. This study also highlights the need for sensitive motor coordination assays and automated analyses to detect early behavioral markers of PD. Our findings suggest the rotenone exposure environmental mouse model as a platform for studying prodromal PD, which may facilitate the development of early detection and intervention strategies.

## Methods

### Mouse housing and care

All experimental procedures were reviewed and approved by the MIT Committee on Animal Care (under protocol #2605000928) and performed according to the guidelines set by National Research Council’s Guide for the Care and Use of Laboratory Animals and the U.S. Public Health Service Policy on Humane Care and Use of Laboratory Animals. Seven-months old male C57BL/6 mice were obtained from the Jackson Laboratory. Up to 5 mice were housed in individual cages under standard laboratory conditions. Mice were kept in a 12:12 reverse light cycle and provided standard food and water *ad libitum*. Functional assays were performed during the dark cycle and at the same time of the day to account for possible circadian rhythm influences on GI function^53^.

### Rotenone preparation and dosing

Rotenone (Sigma-Aldrich, R8875) was first prepared in 50X stock solution that was diluted in 100% dimethyl sulfoxide (DMSO) (Sigma-Aldrich, 472301). The rotenone solution was then and diluted in medium-chain triglyceride, Miglyol 812N (IOI Oleochemical, Germany) to obtain a final working concentration of 2.5mg/kg rotenone in 98% Miglyol 812N and 2% DMSO. The rotenone was prepared fresh every day for 5 days and stored at room temperature shielded from light. Mice received once-daily intraperitoneal injections of rotenone (2.5mg/kg i.p.) or vehicle (98% Miglyol 812N, 2% DMSO) for 5 consecutive days.

### Behavioral assessments

#### Open Field Test (OFT)

Locomotor activity and anxiety-like behavior were measured using the OFT. Animals were habituated to the open-field chamber for 2 days before each test day. Each mouse was placed in the center of the 30 x 30 cm open-field chamber and were recorded for 20 min before being returned to their home cage. Multiple behavioral parameters were obtained and analyzed using EthoVision Software (EthoVision XT, Noldus Information Technology, Inc.) including total distance travelled, time spent in periphery and center, and velocity.

#### Rotarod test

Motor coordination, balance and motor learning were evaluated using the rotarod test. Mice were placed on the rotarod apparatus (Five lane rota-rod for mouse, ENV-574M, Ugo Basile, Italy) which was accelerated from 4 to 40 rpm for a 5-min period. Mice underwent habituation for 2 days on constant-speed loading mode before being tested on 2 consecutive days using the accelerating (4 to 40 rpm) protocol. The time taken for each mouse to fall from the rod (latency) was recorded. The average latency to fall was used for analysis.

#### Pole test

The pole test was used to assess bradykinesia and motor coordination. The mice were acclimated to the pole test set-up for 2 days before test day. One mouse at a time was placed head facing upward at the top of a vertical pole. The mouse was then allowed to turn downward and descend into the home cage positioned at the base. The time taken for a mouse to turn downwards (time taken to turn 180°) and the time taken to descend the pole were recorded. A maximum cut-off of 60s was ensured for the descent. Each mouse completed 3 trials, and the average of the trials was calculated.

#### Postural Instability Test (PIT)

The PIT was used to evaluate forelimb motor function and postural control. In prior literature, measurement of minimum forward displacement to trigger a compensatory step in the PIT was primarily done in rats and quantified in a manual fashion by tracking the position of the nose during the forward step as a proxy for postural correction^26,54^. Since mice are smaller in size and more prone to lateral head movement during the test, nose displacement was a less stable metric, and manual measurement was more subjective due to steps taken at a smaller scale. Therefore, we adapted the test and developed an automated analysis pipeline based on computer-vision methods to measure the displacement based on the distance between successive forelimb placements (step size). A transparent plexiglass board (marked from 0 to 10 cm using straight lines to indicate every cm) was placed on top of a 30 x 30 cm acrylic chamber which had a camera fixed in it to monitor the paw movement of the mice from beneath. Mice were held vertically with the head facing downward. The left forelimb was gently restrained against the torso while the contralateral forelimb was allowed to contact the surface. The front right paw of the mouse was placed at the 0 cm mark of the plexiglass board and then moved forward slowly until a compensatory step was initiated. Each mouse underwent 3 trials, and mean displacement was calculated for analysis.

#### Automated analysis of PIT

DeepLabCut was used to perform pose estimation and trace movement of mice recorded from beneath the clear plexiglass board positioned to enable clear visualization of the paws using videography at 60 frames per second (**Supplementary Fig. S1a-c)**. The trace was then used to extract the distance to trigger compensatory step which has been previously used as a metric of motor coordination and postural adjustment^55,56^. Specifically, the mid-paw point of the right forelimb being used as the Y-position (**Supplementary Fig. S1d-e**). The DeepLabCut’s output was imported to MATLAB for further analysis where the step sizes were calculated (**Supplementary Fig. S1f**. The number of trials, steps, distance between each step and average distance of each step (mm) was calculated. Step distances were calculated using the absolute difference in Y-positions between consecutive steps, based on the procedure used to manually analyze the PIT (**Supplementary Fig. S1g**). The validity of the automated analysis was established by manually labelling videos of the postural instability test from one of the timepoints (n=10, 5 rotenone and 5 vehicle) and comparing manual and automated results of step distance. The mean step distance and error bars for each scoring method was calculated (**Supplementary Fig. S1h)**. The intraclass correlation coefficient, which quantifies the degree of correlation and agreement between measurements, was 0.79. This value exceeds the 0.75 threshold defined by Koo and Li ^57^, which indicates good reliability and agreement between the manual and automated analysis. Moreover, the DeepLabCut analysis demonstrated a 93% accuracy compared to manual scoring and reduced the analysis time from 5-15 minutes per video to 2-4 minutes per video. These results suggest that the DeepLabCut can reliably assist analysis with a high degree of accuracy and improved efficiency.

#### Elevated Plus Maze (EPM)

Anxiety-like behavior was assessed using EPM. The EPM apparatus consisted of four arms (30 cm in length, 5 cm in width) with two open and two enclosed arms with opaque walls (15 cm in height) arranged in a plus configuration connected by a central area (6 cm in length, 6 cm in width). This setup was elevated 40 cm above the floor, and the testing room was maintained under dim, diffuse lighting to avoid excessive aversive stimulation. The mice were placed in the center and allowed to explore the maze for 5 min. Behavior was recorded using an overhead camera and analyzed using EthoVision Software (EthoVision XT, Noldus Information Technology, Inc.). The time spent in open, closed arms and in center were measured.

#### Catwalk gait assay

The catwalk gait assay was used to measure fine gait parameters. Mice were placed inside a column (7.5 cm in width, 8 cm in height, 150 cm in length) with a transparent bottom, and a high-speed camera placed underneath recording the paw movements at 60 frames per second. Mice traversed the corridor voluntarily from end to another. The feet, nose and tail positions were tracked across frames using motion capture software and low pass filtered using a 6^th^ order double pass Butterworth filter. Only uninterrupted crossings with consistent forward locomotion were included for analysis.

### *Ex-vivo* colonic motility assay

Mice were anesthetized with 3-5% isofluorane (1-2L oxygen) and euthanized by cervical dislocation. Krebs buffer was prepared by mixing 1L Krebs-Henseleit Buffer (Sigma-Aldrich, K3753) in 1L of distilled water, 0.373g of calcium chloride dihydrate (AA Blocks, AA0001M2), 2.1g of sodium bicarbonate (Sigma-Aldrich, S6014). The entire colon, from cecum to rectum, was rapidly excised and transferred to ice-cold (4°C) Krebs-Henseleit Buffer solution continuously bubbled with carbogen (95% Oxygen; 5% CO2, Linde, OX CD5C-T). Surrounding mesenteric tissue was carefully removed under a dissecting microscope. Luminal contents were gently flushed using a blunt needle attached to a 5ml syringe filled with chilled Krebs buffer. The cleaned colon was mounted in a temperature-controlled organ bath chamber infused with oxygenated Krebs solution maintained at 35°C with a flow rate of 3.5mL/min. The bath solution was continuously aerated with carbogen throughout the experiment and preparations were allowed to equilibrate for 30 min prior to data acquisition. Following equilibration, colonic motor activity was recorded for 15 min using high-resolution videos using Gastrointestinal Motility Monitor software (GIMM; Catamount Research and Development, St. Albans, VT, USA).

### Whole gut transit time

Whole gut transit time was determined using previously described carmine dye assay^58^. Mice were first fasted 1 hour prior to the study. The mice were then administered 150 uL of 6% carmine dye (Sigma-Aldrich, C1022) in 0.5% methyl cellulose solution (Sigma-Aldrich, M0512) in distilled water in a red-light illuminated room by oral gavage. The mice were then placed in individual cages with *ad libitum* access to water and food pellets. The cages were checked for the presence of a red-dyed fecal pellet briefly every 15 min. Transit velocity for the colon was approximated by dividing the colonic length by the whole gut transit time.

### Tissue preparation

Mice were deeply anesthetized and transcardially perfused with chilled phosphate-buffered saline (PBS) (Corning®, 21-040-CV), followed by 4% paraformaldehyde (PFA) which is diluted from 32% PFA (Electron Microscope Sciences 15714-S) in 1X PBS. Tissues were dissected, post-fixed in 4% PFA at 4°C for 24 h and processed for paraffin embedding or cryoprotection in 30% sucrose (Sigma-Aldrich, S7903), diluted in 1X PBS for frozen sectioning. For paraffin processing, tissues were dehydrated through graded (70-100% diluted in distilled water) ethanol washes, cleared in xylene (Sigma-Aldrich, 534056) and embedded in paraffin. Sections (4 µm) were cut using a rotary microtome and mounted on charged glass slides. For cryosections, tissues were embedded in O.C.T compound (Tissue-Tek®, 4583), frozen and sectioned at 10 µm using a cryostat.

### Histology

#### Hemoxylin & Eosin (H&E)

Sections were deparaffinized in xylene and rehydrated through graded ethanol (100 – 50%, diluted in distilled water) to deionized water. Slides were stained with hematoxylin, rinsed in running tap water, differentiated as required, and counterstained with eosin. Sections were cover-slipped with Fluoromount-G® mounting media (Southern Biotech, 0100-01). H&E staining was used for general morphological assessment.

#### Alcian Blue

For detection of acidic mucopolysaccharides and goblet cells, sections were deparaffinized and rehydrated to water. Slides were incubated in Alcian Blue solution (pH 2.5) to visualize acidic mucins, rinsed, and counterstained with Hematoxylin. Sections were dehydrated, cleared, and mounted for imaging.

### Immunohistochemistry

#### Avidin-Biotin Complex (ABC)

The sections underwent antigen retrieval (heat-mediated) followed by quenching of endogenous peroxidase activity using hydrogen peroxide (Sigma-Aldrich, 216763). Non-specific binding was blocked using normal donkey serum (Jackson ImmunoResearch, 005-000-121), hereafter referred to as blocking buffer. Sections were incubated overnight at 4°C with tyrosine hydroxylase (TH) primary antibody, rabbit-anti-tyrosine hydroxylase antibody (1:1000, Novus Biologicals NB300-109), diluted in blocking buffer. After washing, biotinylated secondary antibody (1:200, goat anti-rabbit IgG) was applied, followed by incubation with Vectastain Elite ABC reagent (Vector Laboratories, PK-6101). Signal was visualized using 3,4’-diaminobenzidine (DAB) substrate (Thermo Scientific, 34065), producing a brown precipitate. Sections were counterstained with hematoxylin, dehydrated, cleared and protected with cover glass. Negative controls included omission of primary antibody.

#### Immunofluorescence

Cryosections were rinsed in PBS, permeabilized and blocked in 5% normal donkey serum containing 0.3% Triton X-100 (Sigma-Aldrich, X100) for 1 h at room temperature. Primary antibodies were applied in 5% normal donkey serum overnight at 4 °C. After 3 PBS washes, fluorophore-conjugated secondary antibodies were applied for 1-2 h at room temperature in the dark. Nuclei were counterstained with DAPI (Thermo Scientific, 62248). Sections were mounted with Fluoromount-G® mounting medium (Southern Biotech, 0100-01). Images were acquired using a confocal microscope (Leica Stellariss SP8, Leica Microsystems, Wetzlar, Germany) with consistent exposure settings at 488nm, 633nm wavelengths respectively. Quantification of fluorescence intensity, cell counts, or colocalization was performed using ImageJ^59^ with uniform background subtraction and thresholding. For brain tyrosine hydroxylase (TH) staining, rabbit-anti-tyrosine hydroxylase antibody (1:1000, Novus Biologicals NB300-110SS) and goat anti-rabbit IgG (H + L), Alexa Fluor 633 (1:1,000, Thermo Scientific A-21070) were used as primary and secondary antibodies respectively. For colon TH staining, primary (rabbit anti-TH antibody (1:1000, Novus Biologicals NB300-109), and secondary goat anti-rabbit IgG (H + L), Alexa Fluor 633 (1:1,000, Thermo Scientific A-210701) and Alexa Fluor 488 conjugated rabbit anti-PGP9.5 antibody (1:500, Abcam ab302578) were used. Nuclei were counterstained with 4′,6-diamidino-2-phenylindole (DAPI; 1 µg/mL; Thermo Fisher Scientific, 62248) for 5 minutes at room temperature prior to mounting.

### Statistical analysis

All experimental data were statistically analyzed using GraphPad Prism (version 10.2.0, for Windows, GraphPad Software, Boston, MA, USA), except for gait analysis, which was performed separately in Python. All data are expressed as mean ± standard deviation (s.d.) and alpha level, α = 0.05 was used for all tests. Data normality was assessed using the Shapiro-Wilk test. For comparisons between rotenone-injected and vehicle-injected mice, the Mann-Whitney U test was used for non-normally distributed data. An unpaired two-tailed t-test with Welch’s correction was used for normally distributed data. Longitudinal body weight monitoring over the 9 months were analyzed using repeated-measures two-way analysis of variance (ANOVA) to assess the effects of treatment (rotenone or vehicle) and time. Baseline and endpoint bodyweights were also compared within the rotenone and vehicle groups respectively using the appropriate paired statistical test.

Behavioral assessments conducted at multiple time points (i.e. pole test, OFT, rotarod assay, and PIT) were analyzed using two-way ANOVA followed by Tukey’s multiple-comparisons test. For comparisons of behavioral parameters at single time points (i.e. EPM and catwalk gait assay), an unpaired two-tailed t-test with Welch’s correction was used. For catwalk gait analysis, movement velocity and stride length were compared between rotenone-treated and vehicle-treated mice using the Wilcoxon rank-sum test in Python.

Whole-gut transit time, colonic transit velocity, colon length, TH-positive neuronal counts in the SNpc and colonic myenteric plexus, and colonic α-synuclein immunoreactivity were compared between treatment groups using the Mann-Whitney U test. Quantitative analyses of colonic contraction dynamics, crypt length, and goblet cell and mucin measurements were performed using an unpaired two-tailed t-test with Welch’s correction. All statistical tests were two-tailed, and a p-value < 0.05 was considered statistically significant.

## Acknowledgements

This work was funded in part by the Pappudu Sriram and Dr. Rajesh Venkataramani Parkinson’s Disease Fund at MIT, K. Lisa Yang Brain-Body Center at the Yang-Tan Collective at MIT, McGovern Institute for Brain Research at MIT, and the Director’s Pioneer Award from the National Institutes of Health and National Institute for Complementary and Integrative Health (DP1-AT011991). The funder played no role in study design, data collection, analysis and interpretation of data, or the writing of this manuscript. S.S. is the recipient of the McGovern Institute 25^th^ Anniversary Fellowship. J.M. is a recipient of Harvard-MIT Program in Health Sciences and Technology (HST) Fellowship. K.K.L.P. is a recipient of the Friends of the McGovern Institute Fellowship and Poitras Graduate Student Scholar Fellowship. G.L.E. is a recipient of NSF Graduate Research Fellowship, Mathworks Fellowship, Friends of McGovern Institute Fellowship Y.J.K. is a recipient of the Mathworks Fellowship. R.M. is the recipient of the Friends of McGovern Institute Fellowship and the HST Fellowship.

## Author contributions

S.S. and P.A. designed the study. S.S. performed all in vivo experiments, behavioral assessments, immunohistology and data analysis. K.K.L.P. assisted with behavioral assays. J.M. assisted with colonic motility assay and data analysis. G.L.E. assisted with data analysis of colonic motility assay. Y.J.K. assisted with data analysis of gait assay with the gait analysis pipeline they developed. R.M. assisted with pilot experiments for the rotenone mouse model. R.L. contributed to the figure preparation. All authors contributed to the writing of the manuscript.

## Competing interests

All authors declare no financial or non-financial competing interests.

## Data availability

All data generated or analyzed during this study are included in this published article and its supplementary information file.

## Code availability

The underlying code for fine motor coordination (GaitPattern) was developed in Ref. [33] and is available in Github repository and can be accessed via this link https://gitshare.me/repo/bedb9ac3-ea8a-4200-b352-cdd49e62ce21.

## Notes

https://gitshare.me/repo/bedb9ac3-ea8a-4200-b352-cdd49e62ce21

### Competing Interest Statement

The authors have declared no competing interest.

