## Supplementary Information for "Modelling prodromal Parkinson’s disease: gastrointestinal pathology and early motor deficits in mice after transient rotenone exposure"

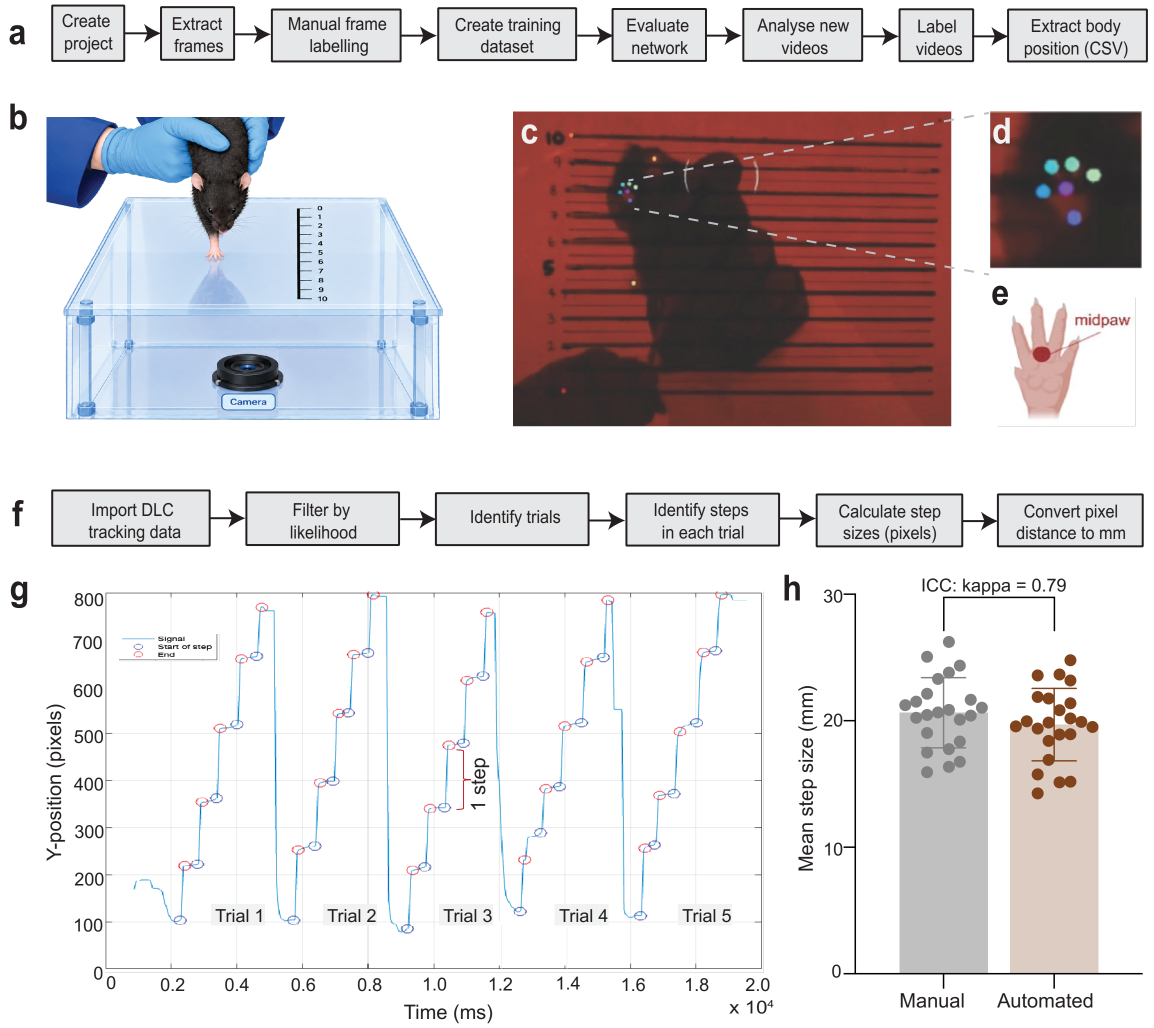


**Supplementary Fig. 1.** Automated postural instability analysis using DeepLabCut. **(a-c)** Analysis pipeline for DeepLabCut and video recording setup from beneath a clear plexiglass board at 60 frames per second for clear visualization of paw positions. **(b)** Illustration of postural instability test was generated by ChatGPT based on prompts given. **(d, e)** Example tracking of the mid-paw point of the right forelimb, used to determine Y-position for step analysis. **(e)** Image of paw generated in BioRender. **(f, g)** Pipeline and analysis for extraction and calculation of step distances, number of steps, and distance between consecutive steps from DeepLabCut output in MATLAB. **(h)** Validation of automated analysis against manual scoring (n = 10 mice; 5 rotenone, 5 vehicle). Data points represent average of step distance calculated from multiple postural stability runs. Mean step distances ± SD are shown. Automated DeepLabCut analysis demonstrated 93% accuracy, reduced analysis time, and good reliability (intraclass correlation coefficient = 0.79).These results demonstrate that DeepLabCut can reliably and efficiently quantify postural adjustments in the postural instability test.
